# Using Patient iPSC-derived Retinal Pigment Epithelial Cells to Evaluate Differential Susceptibility to MEK Inhibitor-Associated Retinopathy

**DOI:** 10.64898/2026.04.11.717944

**Authors:** Lola P. Lozano, Timothy M. Boyce, Andrew P. Groves, Henry L. Keen, Culver Boldt, Robert F. Mullins, Elaine M. Binkley, Budd A. Tucker

## Abstract

**Purpose:** Compare the effect of MEK inhibition on iPSC-derived retinal pigmental epithelial (RPE) cells generated from a patient who developed MEK inhibitor-Associated Retinopathy (MEKAR) versus a patient who did not develop retinopathy.

**Design:** Case-control

**Subjects:** Two female patients with Neurofibromatosis Type 1 who were treated with MEK inhibitors. One patient developed MEKAR, the other did not.

**Methods:** RPE were generated from human induced pluripotent stem cells (hiPSCs) from these two patients. These hiPSC-derived RPE were treated with selumetinib for 10 days.

**Main Outcome Measures:** Phagocytic activity and changes in gene expression

**Results:** As previously reported, there was a significant increase in internalized rhodopsin in phagocytosis assays, yet this was only found in hiPSC-derived RPE from the patient who developed MEKAR. Selumetinib decreased expression of genes related to fluid transport and cell volume, including aquaporins and solute transporters. At baseline, cells from the patients without MEKAR had higher expression of these genes. Interestingly, selumetinib-induced changes in gene expression only reached statistical significance in cells from the patient who did not develop MEKAR, suggesting these changes may be a compensatory protective mechanism. Patients susceptible to forming MEKAR may have increased phagocytosis without a compensatory change in expression of genes related to fluid flux, thereby inhibiting their ability to transport fluid out of the subretinal space.

**Conclusions:** MEK inhibitor-Associated Retinopathy may only affect susceptible patients whose retinal pigment epithelium cannot sufficiently regulate expression of genes related to fluid transport and cell volume, altering the ability of these cells to properly function.

## INTRODUCTION

MEK inhibitors are widely used in oncology to treat many types of malignancies. In particular, these drugs are used long term in patients with Neurofibromatosis Type 1 (NF1) related neurofibromas^1^ Due the ubiquity of the MEK pathway in cell signaling, many patients experience side effects. One side effect that can significantly decrease patients’ quality of life and lead to treatment cessation is MEK inhibitor-associated retinopathy (MEKAR).^2,3^ MEKAR is characterized by subretinal fluid accumulation that leads to decreased visual acuity.^2,4-6^

While the mechanism of MEKAR in unknown, it is hypothesized to cause dysfunction of the retinal pigment epithelium (RPE), preventing these cells from transporting fluid from the subretinal space to the choroid.^6^ In a previous study we demonstrated that inhibition of MEK (via treatment with 10 μM selumetinib) in mature human iPSC-derived RPE cells led to increased phagocytosis and differential expression of genes related to fluid transport and cell volume regulation.^7^ Furthermore, these findings were detectable by as little as 10 days of treatment.

In this study, we compared hiPSC-derived RPE generated from two female patients with NF1 who were both treated systemically with MEK inhibitors yet only one of them developed MEKAR (**Table 1, Figure 1**). To determine if MEK inhibition has a differential effect on hiPSC-derived RPE cells between these patients, we treated cells for ten days and measured changes in phagocytosis and expression of genes relevant to fluid transport.

**Table 1.** Sample demographics. Samples were derived from two female patients with NF1 and treated with the same dosage of MEK inhibitors. However, one patient developed MEKAR and the other did not.

|  | **NF1 with MEKAR** | **NF1 without MEKAR** |
| --- | --- | --- |
| **Sex** | Female | Female |
| **Age at biopsy (years)** | 49 | 21 |
| **NF1 Genotype** | C to T causing premature STOP at chr17.31229061 | G to GT insertion causing frameshift at chr17.31233203 |
| **MEK Inhibitor (dose, frequency, duration)** | Selumetinib<br>50 mg, twice daily<br>5 months at time of biopsy | Trametinib<br>2 mg, once daily<br>Treated for 3 months then discontinued prior to biopsy |

**Figure 1.**
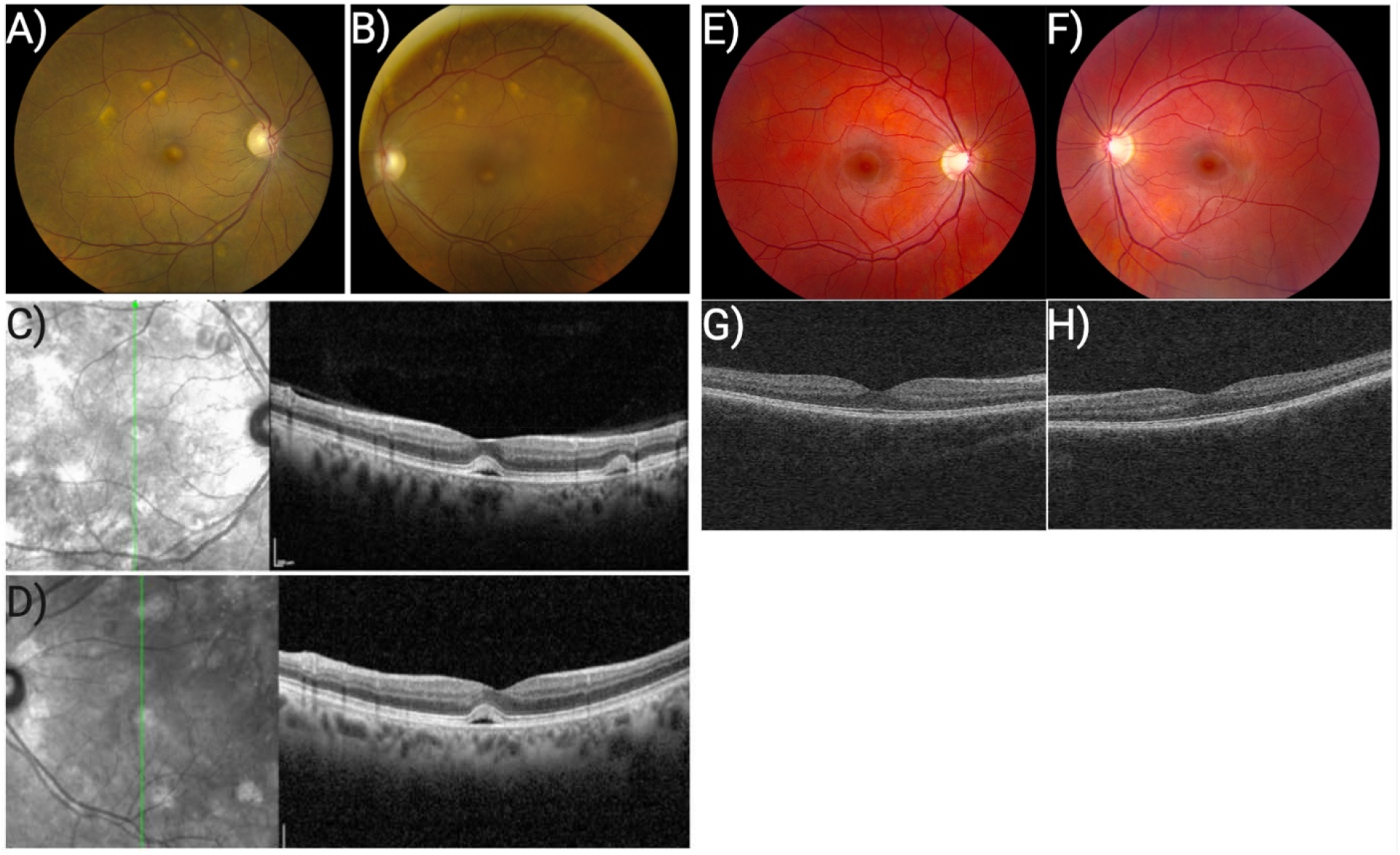
Clinical images of patients treated with MEK inhibitors. A-B) Fundus photos from patient who developed MEKAR of the right (A) and left (B) eye showing subretinal fluid accumulation in the fovea and along the arcades. C-D) OCT images from patient who developed MEKAR showing location of subretinal fluid accumulation in the interdigitation zone between the photoreceptor outer segments and the retinal pigment epithelium in the right (C) and left (D) eye. E-F) Fundus photos from patient who did not develop MEKAR of the right (E) and left (F) eye showing no evidence of subretinal fluid accumulation. G-H) OCT images from patient who did not develop MEKAR of the right (G) and left (H) eyes showing no evidence of subretinal fluid accumulation.

## METHODS

### Study Approval

This study received approval from the University of Iowa Institutional Review Board (IRB No. 200202022). Both patients provided informed consent, and research was conducted in accordance with the Declaration of Helsinki.

### Genetic sequencing

DNA from patient-derived fibroblasts was isolated and submitted for sequencing at the Iowa Institute of Human Genetics – Genomics Division. Reference DNA was submitted from a control female patient without NF1.

High–molecular weight genomic DNA was extracted (Machery-Nagel) and quantified using the Qubit dsDNA HS Assay (Thermo Fisher Scientific). DNA integrity was assessed using a Model 4150 Tapestation (Agilent Technologies). For library preparation, 3–5 ug of genomic DNA was sheared to a target size of ∼15 kb using Covaris g-Tubes (Covaris, Woburn, MA, USA) by centrifugation at 4,700 × g for 60 s, followed by tube inversion and a second centrifugation under identical conditions. Fragment size distribution was confirmed using a Tapestation. Sequencing libraries were prepared using the Oxford Nanopore Technologies (ONT) Ligation Sequencing Kit (SQK-NBD114.24) following the manufacturer’s instructions. Briefly, 1–1.5 ug of sheared DNA underwent DNA repair and end-preparation, followed by native barcode ligation and adapter ligation. Cleanup steps were performed using AMPure XP beads (Beckman Coulter) at a 0.4× bead ratio to retain fragments ≥10 kb. Final libraries were quantified using Qubit and 400–600 ng was loaded per flow cell.

Sequencing was performed on PromethION R10.4.1 flow cells (FLO-PRO114M). Flow cells were primed according to ONT protocols, and libraries were loaded in a total volume of 200 ul. Adaptive sampling was configured in MinKNOW in enrichment mode to target chromosome 17 using a BED file specifying chr17 coordinates (chr17:0– 83,257,441) derived from the GRCh38/hg38 primary assembly reference genome. The hg38 reference FASTA and chromosome 17 BED file were provided to MinKNOW for real-time read classification and selective sequencing. High-accuracy (SUP) base-calling with methylation detection (5mC in CpG context) enabled was performed and unaligned bam/fastq files were generated.

Nextflow (version 21.10.6) was used for running all the bioinformatics workflows mentioned below. The fastq files were first run through the wf-cas9 workflow (version 0.1.8), in which reads were aligned to the human reference genome (hg38) using Minimap2 (version 2.24) and quality control reports were produced. On-target enrichment efficiency was calculated as the proportion of aligned bases mapping to chromosome 17 relative to total aligned bases. After confirming that all samples passed quality control and that adaptive sample yielded significant enrichment, the alignment files were run through the wf-human-variation workflow (version 1.5.2(to identify sequence variants. In this workflow, small variants (i.e. SNPs, indels) were identified using the Clair3 program (version 1.0.1) and large structural variants were identified using the Sniffles2 program (version 2.0.7). The resulting variant files (VCF) were filtered for read depth (DP > 10), and genotype quality (GQ > 10). Annotation of the variants was performed using the online version of the Ensembl Variant Effect Predictor (VEP). Phase analysis of variants was performed using WhatsHap (version 1.2.2). VCF files were phased using the phase command and the haplotag command was subsequently used to add haplotype tags (HP) to sorted bam files. The phased files were then visualized in Integrative Genomics Viewer (IGV, version 2.11.1).

Confirmation of pathogenic mutations identified with long-range sequencing was performed with Sanger sequencing. Amplicons containing each mutation within the *NF1* gene were generated (**Supplementary Table 1**). Primers were designed using PrimerQuest Tool (Integrated DNA Technologies, Inc., Coralville, IA, USA) and prediction for primer specificity verified (UCSC In-Silico PCR). Amplicons were sequenced at the Iowa Institute of Human Genetics – Genomics Division. Results were visualized using SeqMan Ultra (DNASTAR, Inc., Madison, WI, USA).

### Generation of human induced pluripotent stem cells

Human induced pluripotent stem cells (hiPSCs) were generated as previously described from 2 female patients with NF1.^8^ Briefly, fibroblasts were isolated and expanded from a 3 mm dermal biopsy and transduced with CytoTune-iPSC 2.0 Sendai Reprogramming Kit (Invitrogen/Thermo Fisher Scientific, Waltham, MA, USA). hiPSC colonies were isolated and expanded. TaqMan Human Pluripotent Stem Cell Scorecard Panel (Life Technologies/Thermo Fisher Scientific, Waltham, MA, USA) was used to analyze pluripotency and loss of viral transgene expression (**Supplementary Figure 1A-B**). Karyotype analysis was carried out by the University of Iowa Shivanand R. Patil Cytogenetic and Molecular Laboratory following a standard G-banding protocol (**Figure 2A**). Cells were maintained at 37C, 5% CO2, and 20% O2.

**Figure 2.**
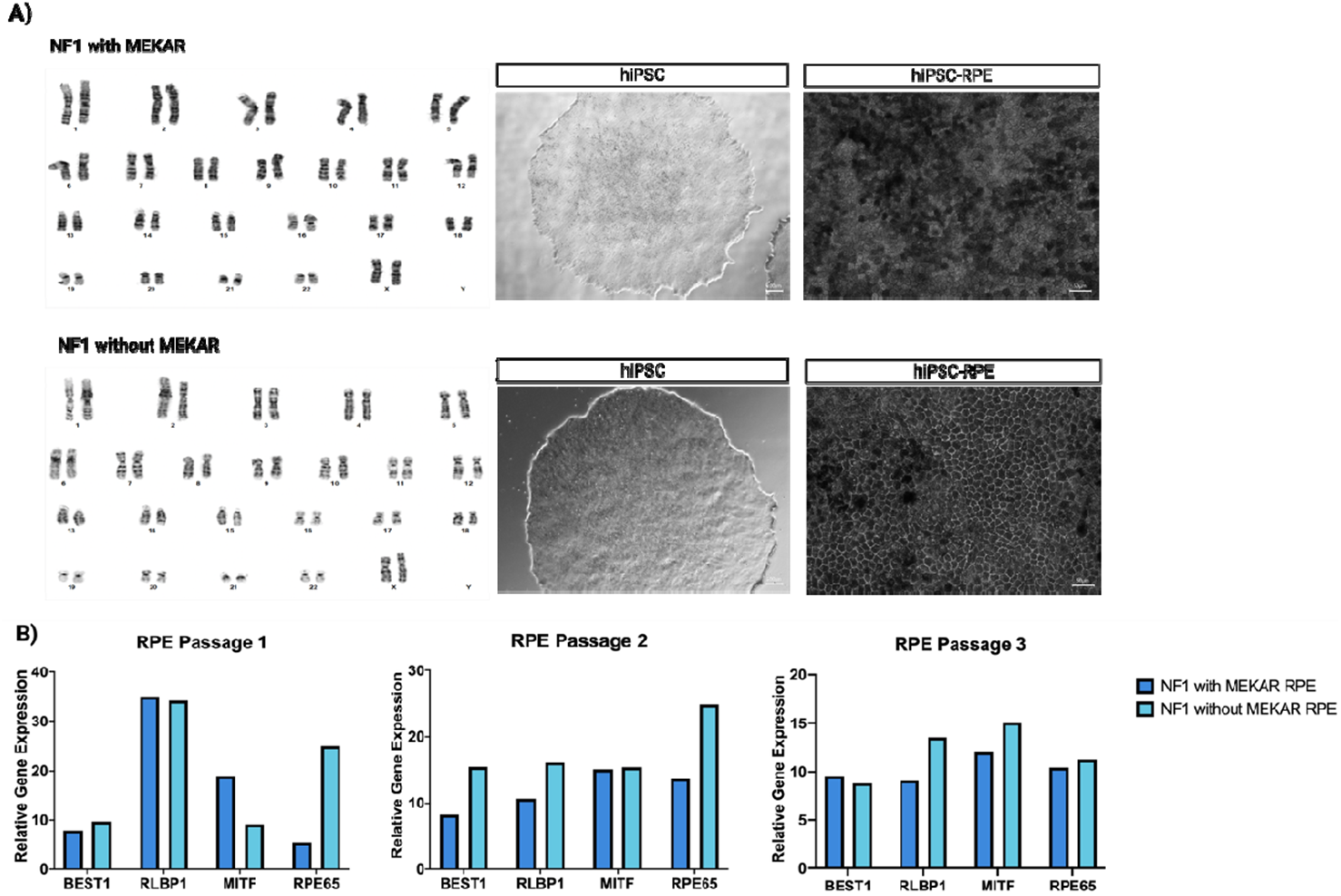
Characterization of patient hiPSC and hiPSC-derived RPE. A) Karyotype G-banding report shows no chromosomal abnormalities for both patient lines used. hiPSCs and hiPSC-derived RPE display expected morphology on phase contrast microscopy. hiPSC-derived RPE sustain expression of RPE-specific gene expression through passages 1-3.

### Differentiation of hiPSC-derived retinal pigment epithelial cells

We followed the rapid, directed differentiation protocol by Foltz and Clegg.^9^ The ROCK-inhibitor Y-27632 (MilliporeSigma, Burlington, MA, USA) was added to cell media at each passage for 4-7 days to increase cell survival and attachment. Briefly, hiPSCs were grown to 80% confluence in Essential 8 Flex media (Gibco/Thermo Fisher Scientific, Waltham, MA, USA) on rhLaminin-521 (Gibco/Thermo Fisher Scientific, Waltham, MA, USA) coated plates and then passaged with ReLeSR (STEMCELL Technologies, Vancouver, BC, Canada) 1:4 onto Matrigel-coated plates and treated for 14 days with growth factors. After RPE expansion, cells were maintained in RPE Supporting Media (RSM), composed of X-VIVO 10 (Lonza, Basal, Switzerland) and 0.2% Primocin (InvivoGen, San Deigo, CA, USA) on tissue culture-treated plates coated with Matrigel (60 ug/ml; Corning Life Science, Tewksbury, MA). Following the ROCK inhibition-dependent extended passage protocol by Croze, Bucholz, Radeke *et al*.,^10^ cells were passaged after reaching confluence every 4-5 days with TypLE Express (Thermo Fisher Scientific, Waltham, MA, USA) and plated at a seeding density of 2.5×10^4^ cells/cm^2^. Only cells at or below passage 3 were used to avoid epithelial-to-mesenchymal transition and loss of RPE morphology (e.g., pigment, cobblestone appearance).^10^ Cells were characterized for cell identity using gene and protein expression (**Figure 2A-B**). Once cells were seeded onto experimental plates at density of 1.0×10^5^ cells/cm^2^ they were allowed to mature for 30 days before drug treatment.

### Pharmacologic Inhibitor of the MAPK/ERK Pathway

Selumetinib (MEK inhibitor) was reconstituted to 10 mM in DMSO (Fisher Chemical, Hampton, NH, USA) per the manufacturer’s recommendations (Sigma-Aldrich, St. Louis, MO, USA). Confirmation of drug efficacy at 10 μM was assessed via Western Blot for ratio of Ph-MAPK to total MAPK (**Supplementary Figure 2**).

### Quantitative Real-Time Polymerase Chain Reaction (RT-qPCR)

Cells were washed twice with 1xDPBS then incubated with 250 ul TrypLE for 5 minutes at 37C. Cells were scraped off and spun down at 10,000 rpm for 5 minutes. Liquid was aspirated and pellets were stored at –20C. cDNA was generated from RNA (200 ng for characterization of cell identity and 231 ng for characterization of differentially expressed genes) using SuperScript IV VILO Master Mix (Thermo Fisher Scientific, Waltham, MA, USA). PCR reactions were run using PrimeTime qPCR probes (Integrated DNA Technologies, Coralville, Iowa, USA) and TaqMan Fast Advanced Master Mix (Thermo Fisher Scientific, Waltham, MA, USA). cDNA was amplified using a QuantStudio 6 Flex Real-Time PCR system (Thermo Fisher Scientific, Waltham, MA, USA). Data were analyzed using delta-delta Ct method in which values were normalized to the *B2M* housekeeping gene and corresponding mRNA values in hiPSCs from the non-NF1 control donor.

### Immunocytochemistry (ICC)

Cells were seeded onto 12-mm round glass coverslips coated in poly-L-lysine (Corning Life Science, Tewksbury, MA) and Matrigel which were placed in 24-well tissue culture treated plates (Corning Life Science, Tewksbury, MA). After 30 days of maturation, cells were washed twice with 1x DPBS then fixed with 4% paraformaldehyde for 20 minutes at room temperature. Washes were repeated followed by blocking for 1 hour at room temperature (**Supplementary Table 2**). Cells were incubated with primary antibody for 2 hours at room temperature. Cells were washed before incubating with secondary antibody and 1:2000 DAPI (Thermo Fisher Scientific, Waltham, MA, USA) for 2 hours at room temperature. Cells were washed then mounted onto microscope slides with coverslips and imaged at 40x using an Olympus BX41 microscope with a digital camera (SPOT Imaging).

### Rod Outer Segment Phagocytosis Assay

Bovine rod outer segments (bROS) (InVision BioResources, Seattle, WA, USA) at a concentration of 10 bROS/cell were added to hiPSC-derived RPE cells in a volume of 500 ul/well of a 24-well plate. Cells were incubated with bROS for 3 hours at 37C and 5% CO2. After 3 hours, all conditions were washed five times with 1xDPBS with calcium and magnesium. Prior to these washes, wells for the Internalized bROS condition were incubated with 2 mM EDTA in PBS for 10 minutes at 37C, 5% CO2 to remove unbound bROS. After washes, Total and Internalized condition wells were collected. Before collecting cells from any condition, 10 uM recombinant human FGF-basic (Peprotech/Thermo Fisher Scientific, Waltham, MA, USA) was added for a 10-minute incubation at 37C, 5% CO2. This step was done to ensure visualization of Ph-MAPK on Western Blot.

### Western Blot Analysis

Cells cultured and treated on 24-well plates were collected then lysed with 75 ul RIPA Lysis and Extraction Buffer with 1:100 Halt Phosphatase and Proteinase Cocktail Inhibitors (Thermo Fisher Scientific, Waltham, MA, USA). Protein concentration was quantified using Pierce BCA Protein Assay (Thermo Fisher Scientific, Waltham, MA, USA) with duplicates of 10 ul per standards and samples. Colorimetric changes in plates were read at 562 nm with the BioTek Cytation 5 microplate reader after 30 minutes of incubation at 37C. 5 ug protein was run on a 4-20% Novex Tris-Glycine gel (Thermo Fisher Scientific, Waltham, MA, USA) for 25 minutes at 225 volts. Broad-range protein transfer to blots was performed using iBlot 3 set to a 6-minute run at 25 volts with low cooling (Thermo Fisher Scientific, Waltham, MA, USA). After incubation with blocking buffer for 1 hour at room temperature or overnight at 4C, blots were probed with primary antibody then washed and probed with HRP-conjugated secondary antibody (**Supplementary Table 3**). Blots were imaged using the iBright™ FL1500 (Thermo Fisher Scientific, Waltham, MA, USA). Blots were then stripped with Restore™ stripping buffer (Thermo Fisher Scientific, Waltham, MA, USA) and probed for beta-actin after imaging to confirm stripping removed previously bound antibody for protein of interest. Densitometry was performed in FIJI ImageJ. All protein bands were normalized to beta-actin.

### Statistical Analysis

Statistical analysis was performed in GraphPad Prism Version 10.2.3 (347) unless otherwise noted. All data were collected from three independent experiments (n=3). Statistical significance was calculated using paired t-test for all RPE cell function assays (alpha = 0.1).

## RESULTS

### Identification of pathogenic mutations

As the NF1 gene is large and identifying pathogenic mutations with gold-standard methods such as Sanger sequencing is often challenging, most patients with NF1 are diagnosed based on clinical phenotype only. However, identifying pathogenic mutations can help better understand disease genotype-phenotype correlations and enable precise genetic manipulations as needed. To screen *NF1* for pathogenic mutations in our two affected patients, we used Oxford Nanopore long-range sequencing. *NF1* variants identified via long-range sequencing were subsequently confirmed using Sanger sequencing. As demonstrated in **Table 1**, the NF1 patient with MEKAR had a premature STOP mutation (C>T) and the NF1 patient without MEKAR had a frameshift mutation (G>GT).

### Impact of MEK inhibition on rod outer segment phagocytosis

To determine if MEK inhibition via selumetinib impacted the ability of iPSC-derived RPE to perform phagocytosis, we added bovine rod outer segments (bROS) to cell media. Three hours after administration, we washed and collected the cells. Protein was isolated and used for Western blot to quantify the fraction of total (membrane bound and internalized) and internalized rhodopsin (RHO) as measure of bROS phagocytosis (**Figure 3C**). There was no difference in total RHO between untreated and treated conditions for each patient (1.9 vs 2.2 for patient with MEKAR, 1.7 vs 1.6 for patient without MEKAR) (**Figure 3A**). There was, however, a significant increase in internalized RHO in treated cells from the patient with MEKAR (1.2 vs 1.9) and no significant difference in the patient without MEKAR (1.4 vs 1.5) (**Figure 3B**).

**Figure 3.**
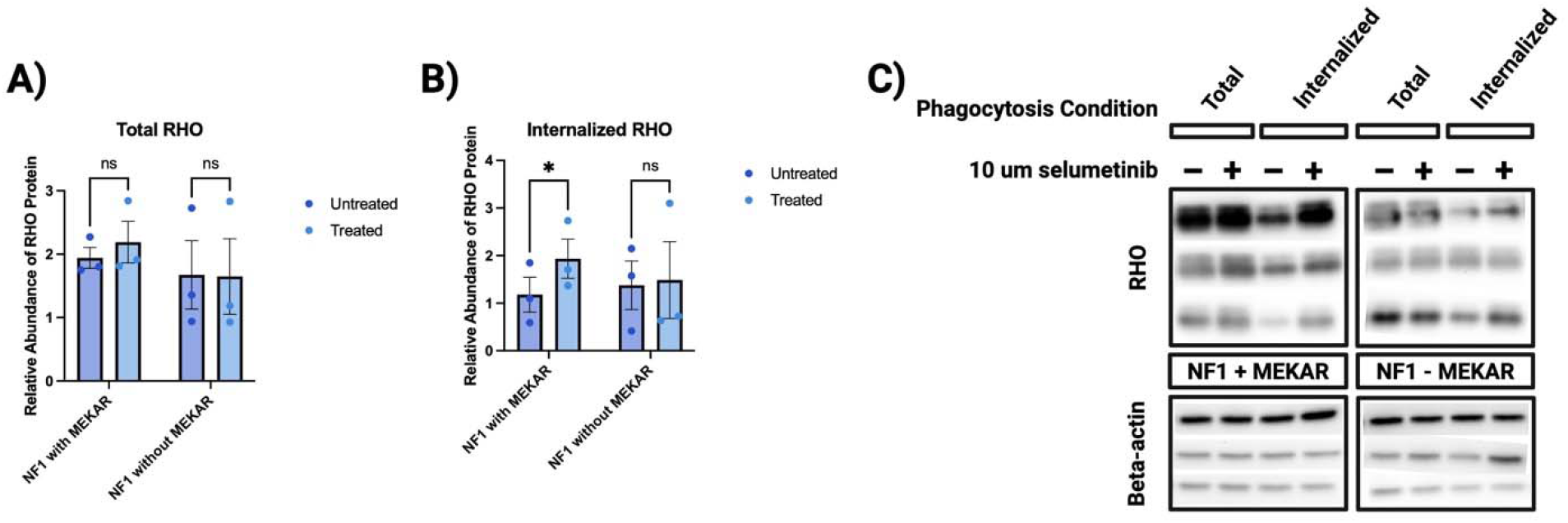
Selumetinib increases rod outer segment phagocytosis in hiPSC-derived RPE of an NF1 patient with MEKAR. A) There was no significant difference in total RHO between untreated and treated conditions in either patient. B) There was a significant increase in internalized RHO in treated cells from the NF1 patient with MEKAR. C) Phagocytosis was determined by Western Blot using antibodies targeted against RHO, the rod-specific photopigment that is absent in RPE cells.

### Differential gene expression between MEKAR and non-MEKAR iPSC-derived RPE cells

To evaluate changes in expression of genes involved in fluid transport and cell volume regulation, which we previously found were dysregulated in iPSC-derived RPE cells treated with selumetinib^7^, a quantitative RT-PCR panel was performed. We found that selumetinib decreased expression of *AQP1, AQP11, ATP6V1C2, LRRC8C, SLC6A6, SLC12A2, SLC16A1*, and *SLC22A23* in both patient lines (**Figure 4A**). Interestingly, expression of AQP3 and AQP7 were unaffected by MEK inhibition. When iPSC-derived RPE generated from the MEKAR patient were compared to the non-MEKAR patient, with the exception of *AQP11*, we found that all of the genes evaluated were expressed at higher levels in non-MEKAR iPSC-derived RPE (**Figure 4B**). Differences in expression were found to be statistically significant for the genes *AQP1, AQP3, AQP7*, and *CLCN5*.

**Figure 4.**
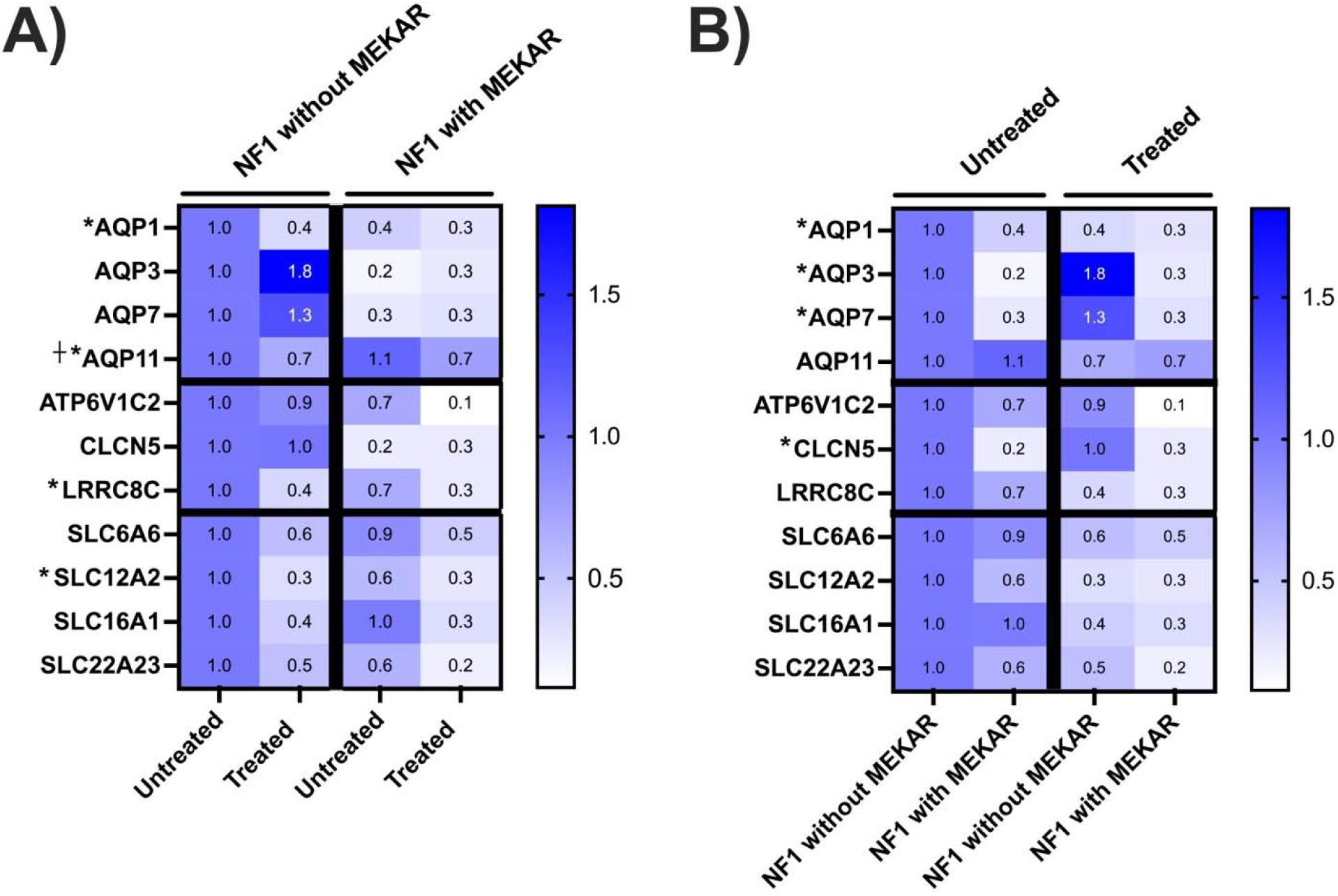
Gene expression changes in hiPSC-RPE treated with MEK inhibition. Selumetinib alters expression of genes related to fluid transport primarily in cells from an NF1 patient without MEKAR. *; significant difference between Untreated and Treated in NF1 without MEKAR. †; significant difference between Untreated and Treated in NF1 with MEKAR. B) The same data rearranged to highlight the expression of genes regulating fluid transport and cell volume regulation are higher in cells from an NF1 patient without MEKAR compared to cells from an NF1 patient with MEKAR. *; significant difference between NF1 with MEKAR and NF1 without MEKAR in Untreated.

## DISCUSSION

The present study contributes to our understanding of the pathophysiology underlying MEKAR by testing a mechanistic hypothesis directly in a human, patient-derived RPE system: that susceptibility to MEKAR reflects differential vulnerability of the retinal pigment epithelium to MEK inhibition^6^. This is clinically and anatomically plausible because the RPE is the key epithelial layer responsible for maintaining outer retinal adhesion and preventing subretinal fluid accumulation via tightly regulated ion and water transport^11-13^.The most direct functional divergence observed between donor lines was seen in the phagocytosis assay: specifically, while selumetinib exposure produced a statistically significant increase in internalized rhodopsin in iPSC-derived RPE cells derived from the MEKAR patient (1.9 vs 1.2), which was consistent with our previous findings^7^, iPSC-derived RPE from the non-MEKAR patient showed no significant change (1.5 vs 1.4). Internalized rhodopsin is metabolized by RPE to ketones which are recycled back into the subretinal space^12^. The polarity of these anions attract sodium and water^14^. Thus, MEK inhibitor induced increases in internalization of outer segments *in vivo* may lead to subretinal fluid accumulation and the MEKAR phenotype.

The present study also found that selumetinib recapitulates directional changes in a focused panel of genes implicated in fluid transport and cell volume regulation, but the pattern of statistical significance differs markedly between lines: the non-MEKAR line shows significant decreases across multiple genes, whereas the MEKAR line shows limited significant change (*AQP11* only)^7^. These findings may suggest that the non-MEKAR line can mount a regulated transcriptional response to MEK inhibition, whereas the MEKAR line is comparatively “transcriptionally inflexible” and instead expresses vulnerability through dysregulated cellular handling of photoreceptor outer segment material.

NF1 is a rare inherited cancer predisposition syndrome, making it challenging to obtain tissue samples for research. We are fortunate to have received biopsies from two patients; however, our sample size is limited to a one-to-one comparison. Additionally, while hiPSC-derived RPE enable *in vitro* experiments dissecting the effect of MEK inhibitors on a single subtype important for retinal health, the isolated nature of this experimental design prevents conclusions about cell-cell interaction and the broader biological environment that likely play a role in MEKAR pathophysiology. Future work using microphysiologic systems and *in vivo* work is needed to fully understand the mechanism underlying this retinopathy.

Many oncology patients treated with MEK inhibitors suffer vision loss due to MEK inhibitor-associated retinopathy (MEKAR). Patients who develop MEKAR may have disrupted function of the retinal pigment epithelium (RPE). In response to treatment with the clinically used MEK inhibitor, selumetinib, human induced pluripotent stem cell-derived RPE cells from a patient with MEKAR had increased phagocytosis and decreased expression of genes related to fluid transport compared to a patient who did not develop MEKAR with treatment. Based on these results, we hypothesize that patients who develop MEKAR have an impaired ability to dynamically regulate expression of genes involved in fluid transport when MEK signaling is suppressed— leading to transient failure of outer retina–RPE fluid homeostasis and resultant subretinal fluid. Alternatively, patients who do not develop MEKAR may simply express levels of fluid transporters such as aquaporins and solute carriers that are sufficient to prevent subretinal fluid accumulation despite MEK inhibition induced downregulation. Future work is needed to confirm these findings *in vivo* and develop appropriate therapies.

## DATA AVAILABILITY

Data are available upon request.

## ACKNOWLEDGEMENTS

Data presented herein were obtained at the Genomics and Bioinformatics Divisions of the Iowa Institute of Human Genetics (RRID: SCR_023422) which is supported, in part, by the University of Iowa Carver College of Medicine. We would like to thank the directors of each of these divisions for their guidance, Kevin Knudtson and Michael Chimenti, respectively.

## Declaration of generative AI and AI -assisted technologies in the writing process

During the preparation of this work the author(s) used ChatGPT in order to assist with drafting the text. After using this tool/service, the author(s) reviewed and edited the content as needed and take(s) full responsibility for the content of the publication.

## REFERENCES

1. Ram T, Singh AK, Kumar A, et al. MEK inhibitors in cancer treatment: structural insights, regulation, recent advances and future perspectives. RSC Med Chem. 2023;14(10):1837–1857.

2. Weber ML, Liang MC, Flaherty KT, Heier JS. Subretinal Fluid Associated With MEK Inhibitor Use in the Treatment of Systemic Cancer. JAMA Ophthalmol. 2016;134(8):855–862.

3. Jeng-Miller KW, Miller MA, Heier JS. Ocular Effects of MEK Inhibitor Therapy: Literature Review, Clinical Presentation, and Best Practices for Mitigation. Oncologist. 2024;29(5):e616–e621.

4. Urner-Bloch U, Urner M, Stieger P, et al. Transient MEK inhibitor-associated retinopathy in metastatic melanoma. Annals of Oncology. 2014;25(7):1437–1441.

5. McCannel TA, Chmielowski B, Finn RS, et al. Bilateral Subfoveal Neurosensory Retinal Detachment Associated With MEK Inhibitor Use for Metastatic Cancer. JAMA Ophthalmology. 2014;132(8):1005–1009.

6. Francis JH, Habib LA, Abramson DH, et al. Clinical and Morphologic Characteristics of MEK Inhibitor-Associated Retinopathy: Differences from Central Serous Chorioretinopathy. Ophthalmology. 2017;124(12):1788–1798.

7. Lozano LP, Jennisch M, Jensen R, et al. Modeling MEK inhibitor-Associated Retinopathy in vitro using human induced pluripotent stem cell-derived retinal pigment epithelial cells. bioRxiv. 2025.

8. Wiley LA, Burnight ER, DeLuca AP, et al. cGMP production of patient-specific iPSCs and photoreceptor precursor cells to treat retinal degenerative blindness. Scientific Reports. 2016;6(1):30742.

9. Foltz LP, Clegg DO. Rapid, Directed Differentiation of Retinal Pigment Epithelial Cells from Human Embryonic or Induced Pluripotent Stem Cells. J Vis Exp. 2017(128).

10. Croze RH, Buchholz DE, Radeke MJ, et al. ROCK Inhibition Extends Passage of Pluripotent Stem Cell-Derived Retinal Pigmented Epithelium. Stem Cells Translational Medicine. 2014;3(9):1066–1078.

11. Yang S, Zhou J, Li D. Functions and Diseases of the Retinal Pigment Epithelium. Front Pharmacol. 2021;12:727870.

12. Lakkaraju A, Umapathy A, Tan LX, et al. The cell biology of the retinal pigment epithelium. Progress in Retinal and Eye Research. 2020;78:100846.

13. Lehmann GL, Benedicto I, Philp NJ, Rodriguez-Boulan E. Plasma membrane protein polarity and trafficking in RPE cells: past, present and future. Exp Eye Res. 2014;126:5–15.

14. Brutsaert EF. Diabetic Ketoacidosis (DKA). MSD Manual. https://www.msdmanuals.com/professional/endocrine-and-metabolic-disorders/diabetes-mellitus-and-disorders-of-carbohydrate-metabolism/diabetic-ketoacidosis-dka. Published 2023. Updated 2023/10. Accessed 2023.

