## Supplementary material for "Using Patient iPSC-derived Retinal Pigment Epithelial Cells to Evaluate Differential Susceptibility to MEK Inhibitor-Associated Retinopathy": n/a

| **Patient Mutation** | **Amplicon (bp)** | **F primer** | **R primer (antisense)** |
| --- | --- | --- | --- |
| **Premature STOP (C>T)** | **733** | CCCACTTTCTTGCCTCTCTT | AGTCCCACTTTCTCATGGTTAC |
| **Frameshift (G>GT)** | **422** | GGCTGATCGGTTTGAGAGATT | TTCCAGCATTGCAACAACTAAC |

**Supplementary Table 1. Primers and amplicons for confirmatory Sanger sequencing.**

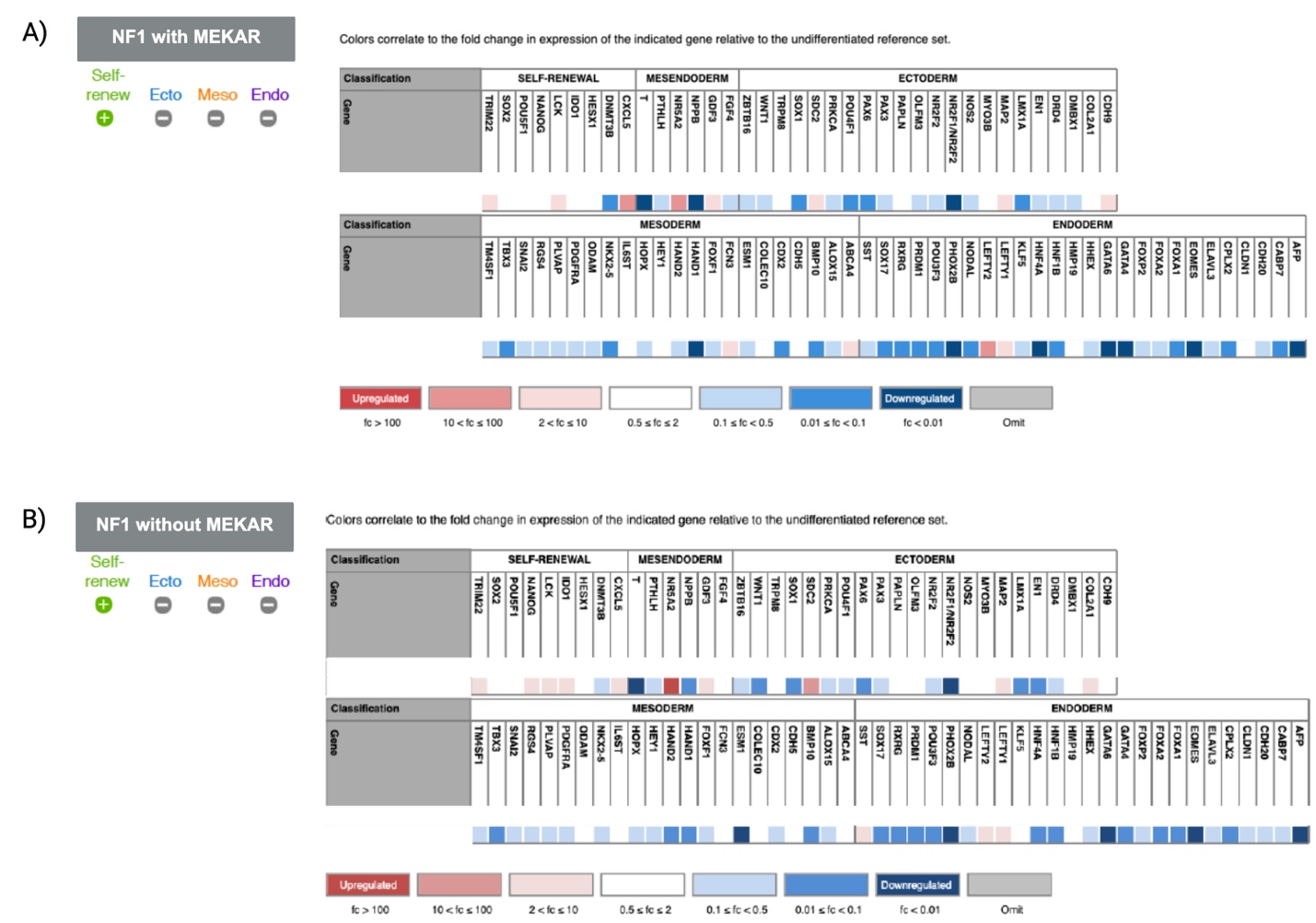

**Supplementary Figure 1. Results of human pluripotent stem cell scorecard panel.** A) hiPSCs from the patient with MEKAR are free of Sendai virus and pluripotent. B) hiPSCs from the patient without MEKAR are free of Sendai virus and pluripotent.

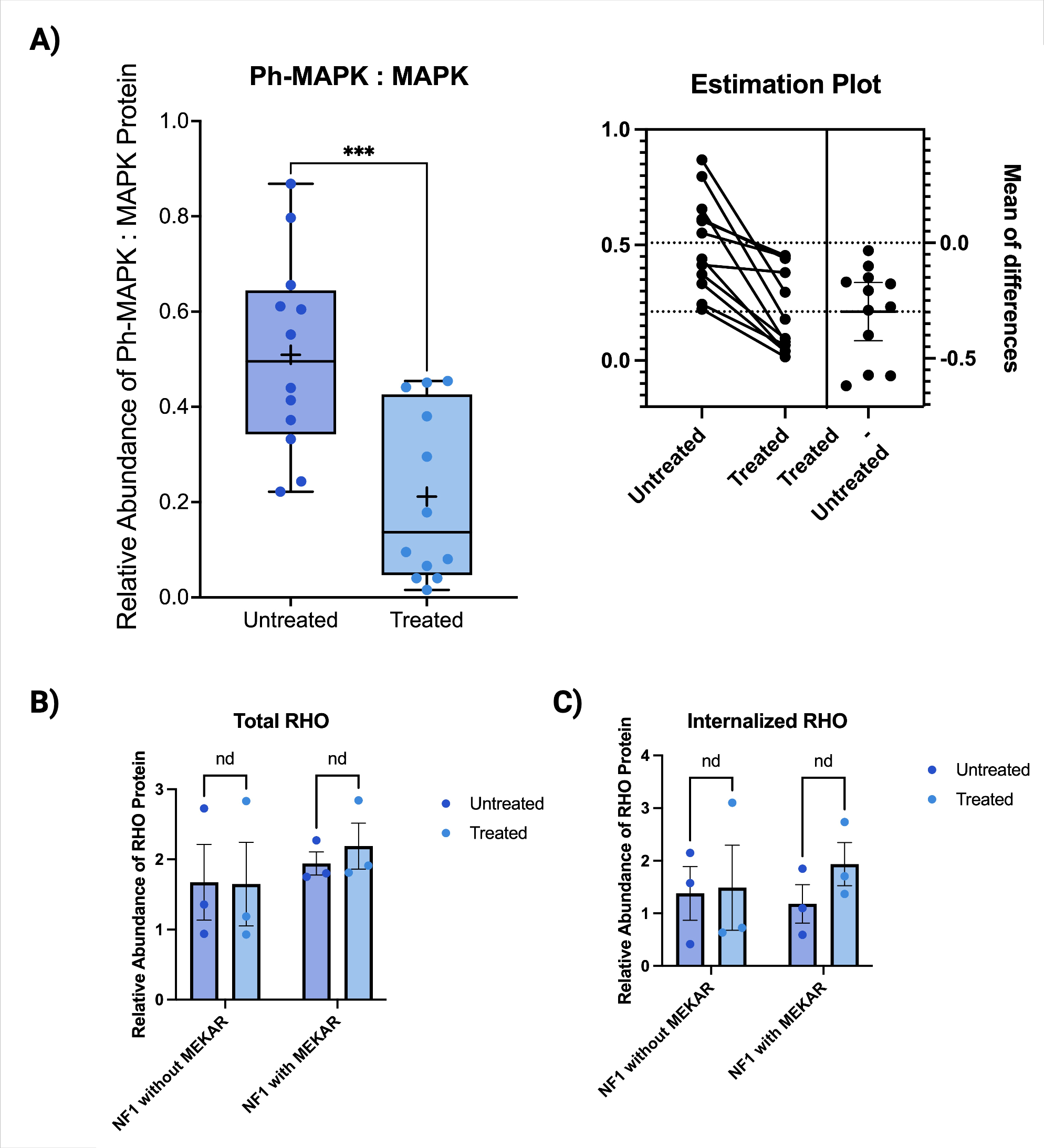

**Supplementary Figure 2. 10 uM selumetinib inhibits phosphorylation and subsequent activation of MAPK.** Paired t-test, p=0.0003.

| **Antibody** | **Company** | **Catalogue #** | **Concentration** |
| --- | --- | --- | --- |
| α-ZO-1 | Invitrogen/Thermo Fisher Scientific | ZO1-1A12 | 1:100 |
| AF647 α-Mouse | ThermoFisher Scientific | A-31571 | 1:1000 |

**Supplementary Table 2 – Antibodies and dilutions for immunocytochemistry.**

| **Antibody** | **Company** | **Catalogue #** | **Concentration** |
| --- | --- | --- | --- |
| α-Beta-actin | Abcam | 8227 | 1:1000 |
| α-MAPK | Cell Signaling Technology | 4696 | 1:1000 |
| α-Ph-MAPK | Cell Signaling Technology | 4370 | 1:1000 |
| α-RHO | MilliporeSigma | MABN15 | 1:10,000 |
| α-Mouse HRP | Cell Signaling Technology | 7076 | 1:2000 |
|  |  |  | 1:50,000 for RHO |
| α-Rabbit | Cell Signaling Technology | 7074 | 1:2000 |

**Supplementary Table 3 – Antibodies and dilutions for western blot.**
